# Cancer Genome Interpreter annotates the biological and clinical relevance of tumor alterations

**DOI:** 10.1101/140475

**Authors:** David Tamborero, Carlota Rubio-Perez, Jordi Deu-Pons, Michael P Schroeder, Ana Vivancos, Ana Rovira, Ignasi Tusquets, Joan Albanell, Jordi Rodon, Josep Tabernero, Carmen de Torres, Rodrigo Dienstmann, Abel Gonzalez-Perez, Nuria Lopez-Bigas

## Abstract

While tumor genome sequencing has become widely available in clinical and research settings, the interpretation of tumor somatic variants remains an important bottleneck. Most of the alterations observed in tumors, including those in well-known cancer genes, are of uncertain significance. Moreover, the information on tumor genomic alterations shaping the response to existing therapies is fragmented across the literature and several specialized resources. Here we present the Cancer Genome Interpreter (http://www.cancergenomeinterpreter.org), an open access tool that we have implemented to annotate genomic alterations and interpret their possible role in tumorigenesis and in the response to anti-cancer therapies.

## New computational tools to support the interpretation of tumor genomes are needed

Cancer is predominantly a genetic disease, caused by the accumulation of so-called “driver” genomic alterations that confer cells tumorigenic capabilities^1^. Thousands of tumor genomes are sequenced every year in research projects and clinical settings around the world. In some cases the whole-genome is sequenced while other focus on the exome or a panel of selected genes. In all cases, the sequencing is followed by the necessity to annotate which of the somatic mutations identified have a possible role in tumorigenesis and treatment response. We call this process ‘the interpretation of cancer genomes’ and it is currently a tedious procedure. One of its major bottlenecks is identifying the driver alterations. A widely employed approach to solve this hurdle consists in focusing on the mutations affecting known cancer genes, i.e., tumor suppressors and oncogenes. These were initially identified through experimentation, giving rise over the past 40 years to a census of human cancer genes^2^. More recently, large re-sequencing projects have provided the opportunity to systematically identify the genes involved in tumorigenesis by detecting signals of positive selection in their alterations pattern across about two dozen malignancies^3–6^. Nevertheless, many somatic variants in tumors, even those in cancer genes, still have uncertain significance and thus it is not clear whether or not they are drivers. Another hurdle in the interpretation of cancer genomes concerns one of its crucial aims: the identification of tumor alterations that may affect treatment options. Unstructured information on the effectiveness of therapies targeting specific cancer drivers is continuously generated by clinical trials and pre-clinical experiments. In summary, novel computational tools are required to address the two aforementioned critical challenges. This includes, on the one hand, methods to estimate the oncogenic effect of the variants observed in a tumor (i.e., identifying validated driver variants and providing some estimation for variants of unknown significance), and on the other, resources that systematically gather the information on biomarkers of drug response and organize them according to distinct use requirements.

## The Cancer Genome Interpreter

Here, we describe the Cancer Genome Interpreter (CGI), a platform that systematizes the interpretation of cancer genomes and makes it automatic. The specific aim of the CGI is to determine which alterations observed in a tumor are more likely to be drivers and identify those that may constitute biomarkers of response to therapies (Fig. 1; details in Supp. Note I). CGI relies on existing knowledge collected from several resources and on computational methods that annotate the alterations in a tumor according to distinct levels of evidence. The tool is a freely available web-resource under an open license, which is intended to facilitate its use by cancer researchers and medical oncologists (http://cancergenomeinterpreter.org). In the following sections we present a blueprint for the interpretation of cancer genomes and describe its implementation in the CGI.

**Figure 1.**
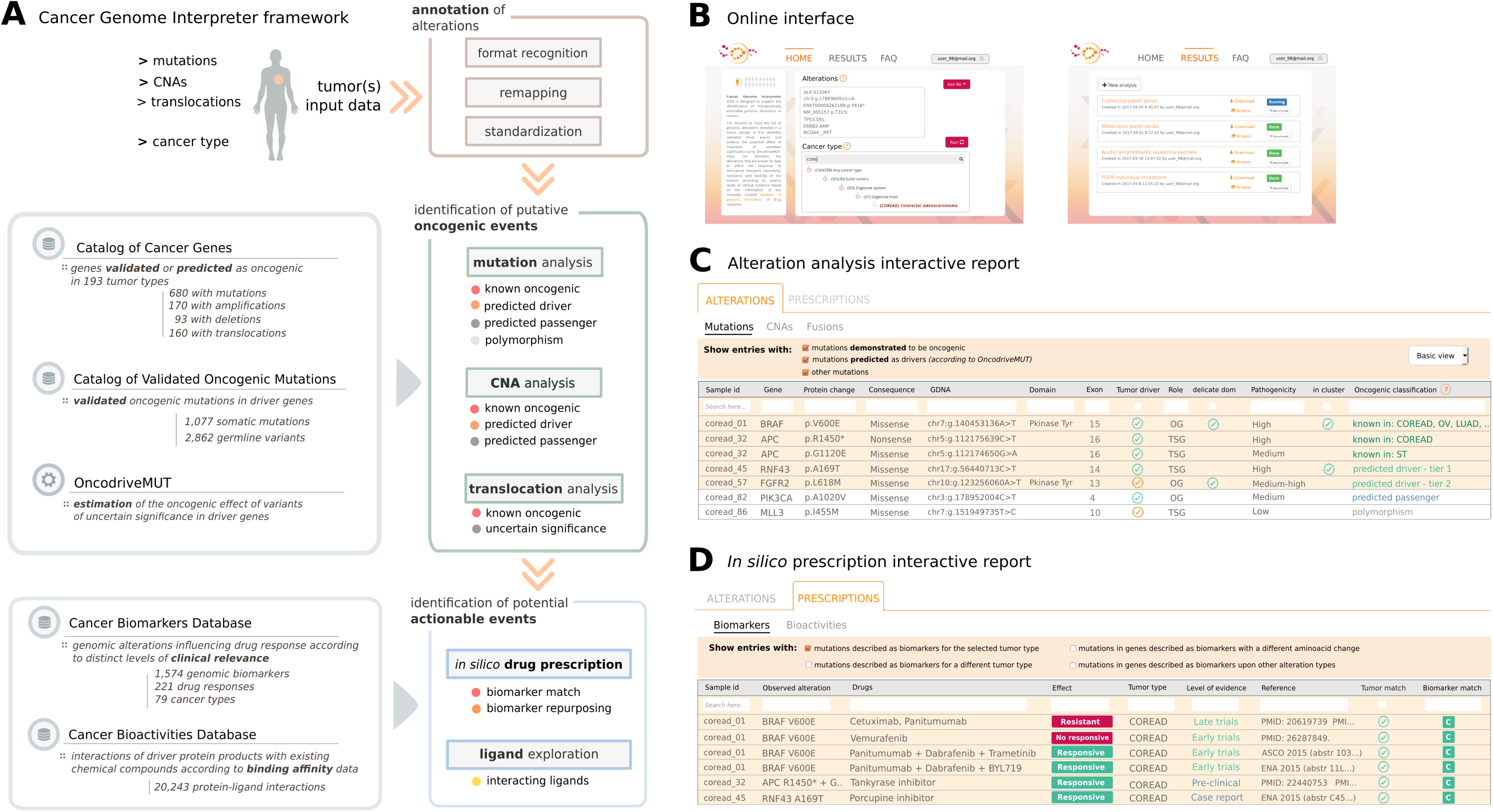
Cancer Genome Interpreter. **(a)** Outline of the CGI workflow. With a list of genomic alterations in a tumor of a given cancer type as input, the CGI automatically recognizes the format, remaps the variants as needed and standardizes the annotation for downstream compatibility. Next, it identifies known driver alterations and annotates and classifies the remaining variants of unknown significance. Finally, alterations that are biomarkers of drug effect are identified. **(b)** The CGI may be run via web at http://cancergenomeinterpreter.com (left panel), or through an API. The web results can be stored in a private repository (right panel) for their management. The results of the CGI are provided via interactive reports: **(c)** Mutation analysis report (example). It contains the annotations of all mutations, which empowers the user’s review, and the labels for those known or predicted to be drivers by OncodriveMUT. **(d)** Biomarkers-match report (example). It contains the putative biomarkers of drug response found in the tumor organized according to distinct levels of clinical relevance. These web reports are interactive and configurable by the user.

## A comprehensive catalog of cancer genes across tumor types

One of the main aims of the interpretation of cancer genomes is to identify the alterations responsible for oncogenic traits. We propose that this process begins with a focus on alterations that affect the genes capable of driving the growth of a particular tumor type. Therefore, we compiled a catalog of genes involved in the onset and progression of different types of cancer, obtained via different methods and from different sources (Supp. Note II). First, we collected genes that have been experimentally or clinically verified to drive tumorigenesis from manually annotated resources^2,7–10^ and the literature. Second, we exploited the bioinformatics results from the analysis of large tumor cohorts re-sequenced by international efforts such as The Cancer Genome Atlas and the International Cancer Genome Consortium^11,12^. On detail, we identified genes whose somatic alterations exhibit signals of positive selection across 6,729 tumors representing 28 types of cancer^4^. In addition, we retrieved the mode of action of each of these cancer genes (i.e., whether they function as an oncogene or a tumor suppressor), curated following state-of-the art knowledge when available and otherwise estimated *in silico*^13^. The resulting Catalog of Cancer Genes currently comprises 837 genes with some evidence of being drivers in 193 different cancer types (Fig. 2a). We annotated each of these genes, identifying (i) the malignancies it drives, organized according to available evidence; (ii) the types of alterations involved (mutations, copy number alterations and/or gene translocations); (iii) the original source(s) reporting it; (iv) the context (germline or somatic) in which these alterations are tumorigenic; and (v) its mode of action as appropriate. The Catalog is available for download through the CGI website (https://www.cancergenomeinterpreter.org/genes).

**Figure 2.**
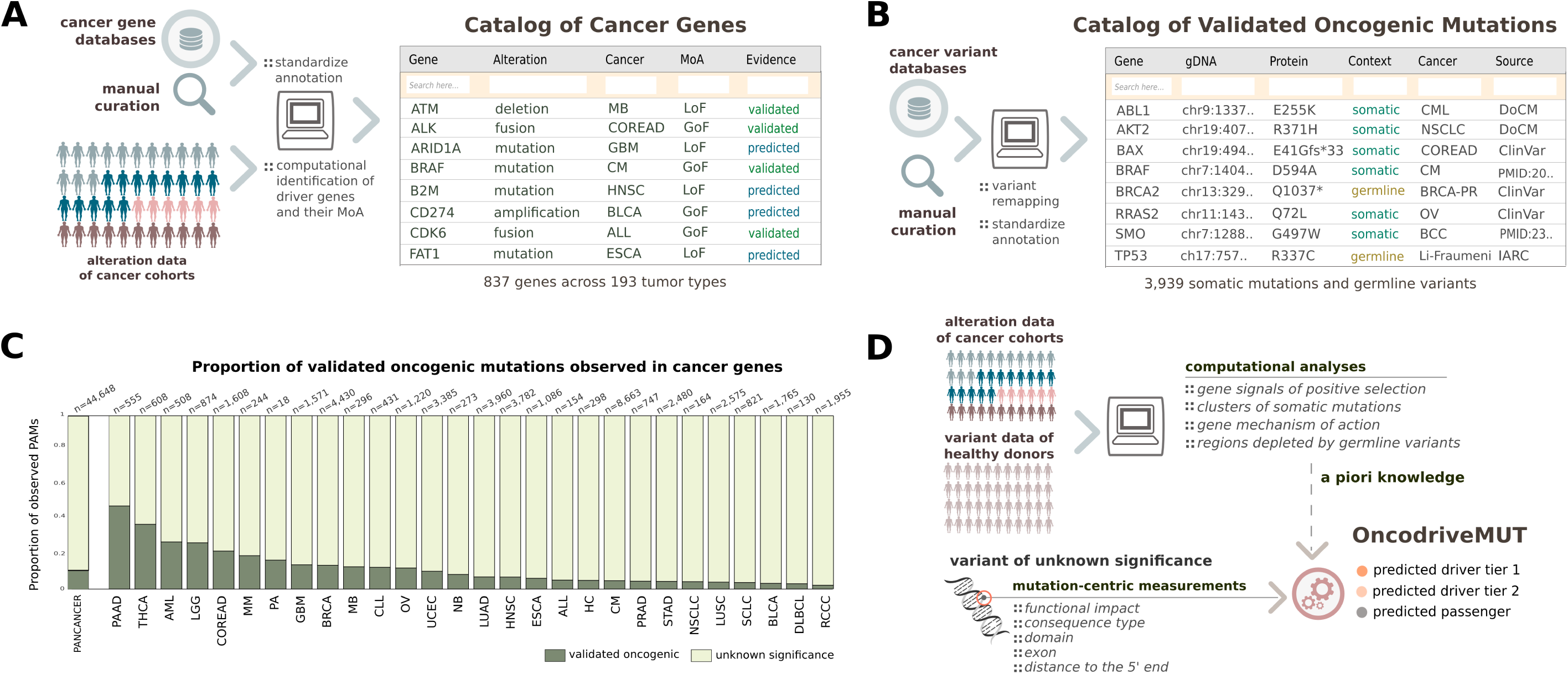
Annotating mutations in cancer genes. **(a)** Catalog of Cancer Genes. Genes that drive tumorigenesis via mutations, copy number alterations and/or translocations are annotated with their mode of action (MoA). **(b)** Catalog of Validated Oncogenic Mutations. Clinically or experimentally validated driver mutations were gathered from manually annotated resources and the cancer literature. **(c)** Proportion of validated mutations observed across the cancer genes of 6,792 tumors. Cancer types acronyms: acute lymphocytic leukemia (ALL); acute myeloid leukemia (AML); bladder carcinoma (BLCA); breast carcinoma (BRCA); chronic lymphocytic leukemia (CLL); cutaneous melanoma (CM); colorectal adenocarcinoma (COREAD); diffuse large B cell lymphoma (DLBC); esophageal carcinoma (ESCA); glioblastoma multiforme (GBM); hepatocarcinoma (HC); head and neck squamous cell carcinoma (HNSC); lower grade glioma (LGG); lung adenocarcinoma (LUAD); lung squamous cell carcinoma (LUSC); medulloblastoma (MB); multiple myeloma (MM); neuroblastoma (NB); non small cell lung carcinoma (NSCLC); serous ovarian adenocarcinoma (OV); pilocytic astrocytoma (PA); pancreas adenocarcinoma (PAAD); prostate adenocarcinoma (PRAD); renal clear cell carcinoma (RCC); small cell lung carcinoma (SCLC); stomach adenocarcinoma (STAD); thyroid carcinoma (THCA) and uterine corpus endometrioid carcinoma (UCEC). **(d)** OncodriveMUT schema to estimate the oncogenic potential of the variants of unknown significance. A set of heuristic rules combines the annotations obtained for a given mutation with the knowledge about the genes (or regions thereof) in which it is observed, as retrieved from the computational analyses of sequenced cohorts.

## Most mutations affecting cancer genes are of uncertain significance

A key aspect of assessing the mutations observed in cancer genes is the tumorigenic potential of each individual variant, as not all of them are necessarily capable of driving tumorigenesis. Therefore, the CGI next focuses on protein affecting mutations (PAMs) that occur in genes of the Catalog of Cancer Genes. Validated tumorigenic mutations may confidently be labeled as drivers when detected in a tumor. We compiled an inventory that currently contains 3,939 such validated driver or cancer predisposing variants from dedicated resources^7–10, 14^ and the literature (Fig. 2B and Supp Note III). This Catalog of Validated Oncogenic Mutations is available for download through the CGI website (https://www.cancergenomeinterpreter.org/mutations). In the pan-cancer cohort of 6,792 sequenced tumors^4^ only 4,142 (630 unique variants) of the 44,648 PAMs found in cancer genes appear in this Catalog. In other words, 90.7% of all PAMs that affect cancer genes in this cohort are currently of uncertain significance for tumorigenesis, a proportion that varies widely per gene and tumor type (Fig. 2c and Supp Note VII). This highlights the need for a means to estimate the tumorigenic potential of these variants. We reasoned that several features of each specific mutation as well as of the genes affected by them could help address this question. Moreover, we propose that some of these features of interest can be extracted from the analyses of large sequenced cohorts of healthy and tumor tissue^4, 15^. Examples of relevant attributes include the following: i) the tumorigenic mode of action of the gene in that cancer (oncogene or tumor suppressor); ii) the consequence type of the mutation (e.g. synonymous, missense or truncating); iii) its position within the transcript; iv) whether it falls in a mutational hotspot or cluster; v) its predicted functional impact; vi) its frequency within the human population; and vii) whether it occurs in a domain of the protein that is depleted of germline variants. The CGI assesses the tumorigenic potential of the variants of unknown significance via OncodriveMUT, a rule-based approach that combines the values of these features (Fig. 1C; Supp. Note IVa). To assess the performance of OncodriveMUT in the task of classifying driver and passenger mutations, we used the Catalog of Validated Oncogenic Mutations (n=3,939) and a collected set of neutral PAMs affecting cancer genes (n=1,247). We found that OncodriveMUT separates the variants of these two data sets with 91% of accuracy (Matthews correlation coefficient, 0.78) (Supp Note IVb). Furthermore, the predictions of OncodriveMUT exhibited a high concordance with the results of experiments assessing the tumorigenic effect of other mutations that are uncommonly seen in cancer^16–19^ (Supp Note IVb). In summary, the CGI annotates the mutations affecting cancer genes with features relevant to their potential role in cancer to facilitate the user’s review, identifying validated drivers and classifying the most likely drivers among the variants of unknown significance.

## A database of genomic determinants of anti-cancer drug response

The second major aim of the effort to interpret cancer genomes is to identify which of the tumor alterations may shape the response to anti-cancer therapies. Findings about the influence of genomic alterations on drug response are continuously generated and reported through publications, clinical trials and conference communications. The challenge resides in gathering relevant results into an easy-to-use resource, and organizing them according to the needs of different users. The CGI employs two resources to explore the associations between gene alterations and drug responses. The first is the Cancer Biomarkers database, an extension of a previous collection of genomic biomarkers of anti-cancer drug response^8^, which currently contains information on 1,574 genomic biomarkers of response (sensitivity, resistance or toxicity) to 221 drugs across 79 types of cancer. Negative results of clinical trials, e.g. the unsuccessful use of BRAF V600 inhibitors as a single therapeutic agent in colorectal cancers bearing that mutation, are also included in the database. Importantly, these biomarkers are organized according to the level of clinical evidence supporting each one, ranging from results of pre-clinical data, case reports, and clinical trials in early (I/II) and late phases (III/IV) to standard-of-care guidelines. The database is under continuous update by a board of medical oncologists and cancer genomics experts (Fig. 3A and Supp. Note V). The second resource is the Cancer Bioactivities database, which currently contains information of 20,243 chemical compounds-protein product interactions that may support novel research applications. We built this database by compiling a catalog of available results from bioactivity assays of small molecules interacting with cancer genes. This information was obtained by querying several external databases (Supp. Note VI). The CGI matches biomarkers or target genes in these databases to alterations observed in tumors. Of note, it reports co-occurring alterations that affect the response to a given treatment. This includes the co-existence of biomarkers of resistance and sensitivity to the same drug, and biomarkers of drug sensitivity that depend upon simultaneous genomic events.

**Figure 3:**
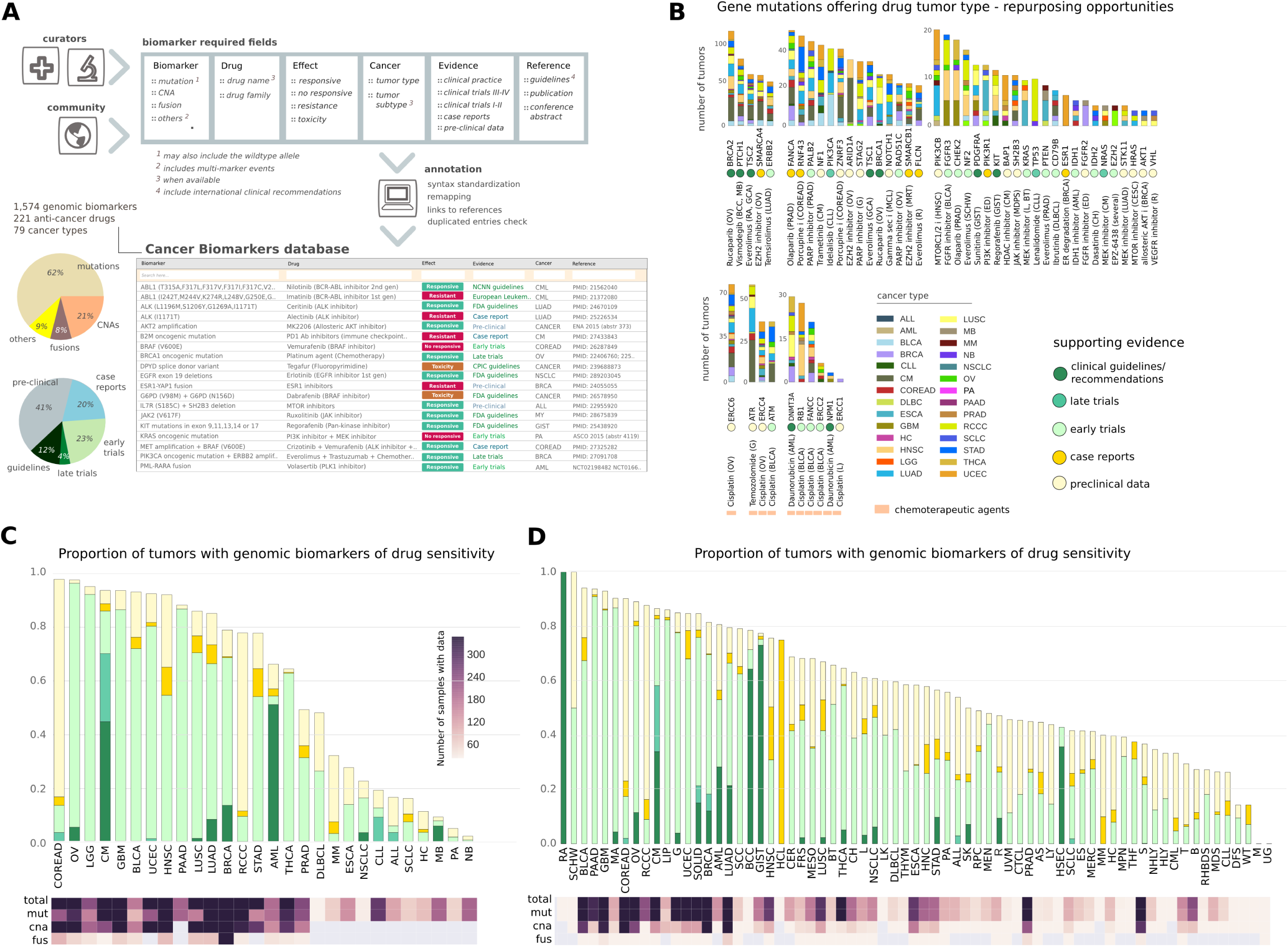
Cancer Biomarkers Database. **(a)** A board of clinical and research experts gather the genomic biomarkers of drug response to be included in the Cancer Biomarkers database through periodic updates. Upper part of the panel displays the simplified schema of the data model. The clinical/research community is encouraged to provide feedback to edit an existing entry or add a novel one by using the comment feature available in the web service. Any suggestion is subsequently evaluated by the scientific team and incorporated as appropriate. A semi-automatic pipeline annotates any novel entry to ensure the consistency of the attributes, including the variant re-mapping from protein to genomic coordinates when necessary. Lower part of the panel displays some of the 1,574 biomarkers that have been collected in the current version of the database, and the left pie charts summarize the content. **(b)** CGI analyses detect putative driver mutations in individual tumors that are rarely observed in the corresponding cancer type. When these variants are known targets of anti-cancer therapies, they may constitute tumor type repurposing opportunities. The graph summarizes some of these potential opportunities detected by the CGI on 6,792 tumors with exome-sequencing data, which are currently unexplored. The barplots display the overall number of tumor samples (separated by cancer type) in which they were observed. The acronym of the cancer type in which the genomic event is demonstrated to confer sensitivity to the drug is shown in parenthesis following the name of the drug, and the clinical evidence of that association is represented through color circles (note that the clinical guidelines/recommendations label refers to FDA-approved or international guidelines). Targeted drugs and chemotherapies are shown separately. Cancer acronyms that are not included in the Figure 2 legend: RA: renal angiomyolipoma; BCC: basal cell carcinoma; GCA: giant cell astrocytoma; G: glioma; MCL: mantle cell lymphoma; MRT: malignant rhabdoid tumor; and R: renal; CH: chollangiocarcinoma. **(c)** Therapeutic landscape of 6,792 tumors with exome-sequencing data. Fraction of tumors with genomic alterations that are biomarkers of drug response in each cancer type. Colors in the bars denote the clinical evidence supporting the effect of biomarkers in that disease (see evidence colors in panel B). Note that the event with evidence closest to the clinic is given priority when several biomarkers of drug response co-occur in the same tumor sample. The lower part of the graph indicates the total number of samples per cancer type, detailing the number of samples in which mutation, copy number alteration (CNA) and/or fusion data was analysed. Cancer acronyms as in Figure 2 legend. **(d)** Same as panel C for a cohort of 17,462 tumors sequenced by targeted panels and gathered by the GENIE project. Tumors were grouped according to the most specific disease subtype available in the patient information. Cancer acronyms that are not included in the Figure 2 legend are detailed in the Suppl. Material.

In summary, these two databases constitute comprehensive repositories of genome-guided therapeutic actionability in cancer according to current supporting evidences. Both resources are available for download through the CGI website (https://www.cancergenomeinterpreter.org/biomarkers, https://www.cancergenomeinterpreter.org/bioactivities). The integration of these two databases with those developed in parallel by other institutions with similar purposes is currently being undertaken within the framework of the Global Alliance for Genomics & Health^20^, described below.

## Current applications and future prospects of the CGI

The CGI (and the databases gathered for its implementation) are under open license, and the resource can be accessed via the web resource and an Application Programming Interface (API; see Supp. Note Ic and Id). The use of the CGI to automatically interpret cancer genomes has broad potential applications, ranging from basic cancer genomics to the translational setting. One feature of the CGI that makes it particularly suitable to different types of applications is its flexibility. The user can input tumor alterations by uploading files following different standards and/or by typing them in a free-text box. The system is prepared to automatically recognize and re-map as necessary different formats, such as genomic, transcript or protein-based coordinates for mutations (Supp. Note Ib). The use of the CGI can help addressing questions raised in different oncology research settings. A newly sequenced group of tumors may be readily interpreted, as exemplified with the pan-cancer cohort presented in this article. The application of the CGI to the mutations profiled across the whole exomes of these tumors delivered a catalog of putative driver alterations across its 28 cancer types (made available through http://www.intogen.org) (Suppl Note VII). The potential of a comprehensive analysis of individual alterations is illustrated by the identification of uncommon events that may be exploited by drug repurposing opportunities (Figure 3B and Supp Note VII). Overall, the CGI identified 5.2% and 3.5% of the samples in the cohort with genomic alterations that are biomarkers of drug sensitivity used in the clinical practice (FDA-approved or international guidelines) or reported in late (phases III-IV) clinical trials, respectively. When considering biomarkers supported by lower levels of clinical relevance, a total of 62% of the tumors exhibited at least one potentially actionable alteration, a number that largely varied across cancer types (Figure 3C and Supp Note VII). However, this cohort mostly includes samples sequenced at diagnosis and thus they may not reflect the type of tumors that are evaluated by molecular oncology boards at present. We also applied the CGI to the sequencing data of 17,642 tumors recently released by the GENIE project, which gathers more advanced cancers profiled by targeted panels^21^. The CGI identified 8% and 6% in that cohort exhibiting biomarkers of drug sensitivity used in clinical practice or reported in late clinical trials, and overall 72% of these tumors exhibited at least one actionable alteration supported by any level of evidence (Figure 3D and Supp Note VII). In addition, the GENIE cohort exhibited more genomic biomarkers of drug resistance, as expected from tumors with a higher proportion of recurrent/relapse patients (Supp Note VII). These analyses provide a comprehensive state-of-the-art snapshot of the putative genomic drivers of cancer and the landscape of genomic guided therapies as it stands today.

On the other hand, the application of the CGI to analyze the results of drug response observed in tumors with different genomic architecture could contribute to the discovery of novel genomic biomarkers of drug sensitivity or resistance. On detail, the distinction between driver and passenger events allows the development of better predictive models^22^. In the clinical setting, application of the CGI to analyze the list of alterations detected in a patient’s tumor could support decision-making in multiple scenarios, especially in cases of variants of unknown significance that may have implications for response to therapy. Early clinical adopters of the CGI used the resource to support the final decision of the most appropriate clinical trial to enroll cancer patients or explore potential drug re-purposing opportunities for pediatric tumors (see Supp. Note VIII).

Crucial to the performance of the CGI are the maintenance and further development of its two distinct types of resources: the repositories of accumulated knowledge and the bioinformatics methods. As new tumor cohorts are re-sequenced and analyzed, our medium-term plans include further development of the catalogs of cancer genes and oncogenic mutations, including both new malignancies and new genomic elements. In particular, the possibility to identify non-coding cancer drivers^23^ from currently generated whole-genome mutation data will open up the opportunity to explore the actionability of non-coding genomic alterations (https://dcc.icgc.org/pcawg). With respect to the aggregation, curation and interpretation of databases of cancer biomarkers and bioactivities, our team follows the standard operating procedures developed under the umbrella of the H2020 MedBioinformatics (http://www.medbioinformatics.eu/) project, thus ensuring the mid-term maintenance of these resources. The feedback from the community is also facilitated through the CGI web interface. Access to this type of cancer data is crucial for the advance of precision medicine, but is highly complex and difficulty for a single institution to comprehensively manage and update. Multiple efforts with similar purposes are currently underway, including My Cancer Genome, https://www.mycancergenome.org; PMKB, https://pmkb.weill.cornell.edu/; PCT, https://pct.mdanderson.org; OncoKB, http://oncokb.org; CIViC, https://civic.genome.wustl.edu; and JAX-CKB https://ckb.jax.org. Within the Global Alliance for Genomics & Health framework^20^, the Variant Interpretation for Cancer Consortium (http://ga4gh.org/#/vicc) was recently launched with the aim to unify the curation efforts of several institutions, including our own. We envision that individual databases will continue to be maintained to fulfill specific needs^24^, but our long-term impact will largely rely, first, on the establishment of international standards for the collection of data relevant to associations between cancer variant-clinical outcome and, second, on our collective success in encouraging the community to share such knowledge.

In summary, the CGI is a versatile platform that automates the steps we propose for the interpretation of cancer genomes, annotating the potential of the alterations detected in human tumors as cancer drivers and their possible effect on treatment response, according to current levels of evidence. The characteristics of the CGI, and the commitment to maintain it as part of a community effort to keep the resource up-to-date with evolving knowledge, allow its establishment as a widely disseminated, easy-to-use tool for both pre-clinical and translational cancer research settings.

## Acknowledgements

This project has received funding from Fundació La Marató de TV3, the European Union’s Horizon 2020 research and innovation programme 2014-2020 under Grant Agreement No 634143, and by the European Research Council (Consolidator Grant 682398). IRB Barcelona is a recipient of a Severo Ochoa Centre of Excellence Award from the Spanish Ministry of Economy and Competitiveness (MINECO) (Government of Spain) and is supported by CERCA (Generalitat de Catalunya). DT is supported by project SAF2015-74072-JIN funded by the Agencia Estatal de Investigacion (AEI) and Fondo Europeo de Desarrollo Regional (FEDER). CR-P is funded by FPI MINECO grant (BES-2013-063354). AG-P is supported by a Ramon y Cajal fellowship (RYC-2013-14554). We appreciate the support provided by Wanding Zhou for the use of the TransVar method and the work of Elaine Lilly in the edition of the text.

